# Variability of an early developmental cell population underlies stochastic laterality defects

**DOI:** 10.1101/2020.07.20.212282

**Authors:** Roberto Moreno-Ayala, Pedro Olivares-Chauvet, Ronny Schäfer, Jan Philipp Junker

## Abstract

Embryonic development seemingly proceeds with almost perfect precision. However, it is largely unknown how much underlying microscopic variability is compatible with normal development. Here, we quantified embryo-to-embryo variability in vertebrate development, by studying cell number variation in the zebrafish endoderm. We noticed that the size of a sub-population of the endoderm, the dorsal forerunner cells (which later form the left-right organizer), exhibits significantly more embryo-to-embryo variation than the rest of the endoderm. We found that, when incubated at elevated temperature, the frequency of left-right laterality defects is increased drastically in embryos with a low number of dorsal forerunner cells. Furthermore, we observed that these fluctuations have a large stochastic component among fish of the same genetic background. Hence, a stochastic variation in early development leads to a remarkably strong macroscopic phenotype. These fluctuations appear to be associated with maternal effects in the specification of the dorsal forerunner cells.

## Introduction

During embryogenesis, cells acquire their identity due to a combination of external signaling cues and internal competence factors. Embryonic development is remarkably robust towards fluctuations of regulatory factors such as morphogen levels (Barkai and Shilo, 2009; Briscoe and Small, 2015) or genetic variation (El-Brolosy et al., 2019). However, developmental buffering of fluctuations is not perfect, and phenotypic variation can even be observed in mutants from isogenic *C. elegans* strains raised under identical environmental conditions due to noisy gene expression and stochastic variation in genetic interaction partners (Burga et al., 2011; Raj et al., 2010).

Differences in the size of progenitor populations may be another important source of phenotypic variability in higher organisms. However, the degree of variability in cellular ontogenies and their potential phenotypic consequences remain largely unknown. Some notable exceptions are: changes in the subdivision of the primordium that gives rise to the head sensory organs lead to fluctuations in ommatidia number at different levels (inter-individual, inter-strain and inter-specific) in the *Drosophila* genus (Gaspar et al., 2020; Ramaekers et al., 2019); sexual dimorphism and left-right asymmetry of ommatidia number in the ant *Temnothorax albipennis* are related to differences in mating and motion behavior, respectively (Hunt et al., 2018). Here, we set out to systematically quantify the degree of inter-individual cell number variation and its phenotypic consequences, using endoderm specification in the early zebrafish embryo as a model system.

The endoderm, which is induced by high levels of Nodal signaling during early development, contributes to the formation of liver, pancreas, intestine, stomach, pharynx and swim bladder (Warga and Nüsslein-Volhard, 1999). The dorsal forerunner cells (DFCs) are a small group of cells, considered a subset of the endoderm (Alexander et al., 1999; Warga and Kane, 2018) (Figure 1A). They are the precursors of the Kupffer’s vesicle (KV) (Melby et al., 1996, Oteíza et al., 2008), the organ that determines left-right asymmetry in the embryo (Essner et al., 2005) (Figure 1B). Since the endoderm is the smallest germ layer during early zebrafish development (Shah et al., 2019), and since relatively large fluctations of DFC numbers between individual embryos were previously reported (Oteíza et al., 2008, Gokey et al., 2015a), we asked if naturally occurring embryo-to-embryo variation in cell numbers in wild-type embryos could have any phenotypic consequences.

**Figure 1.**
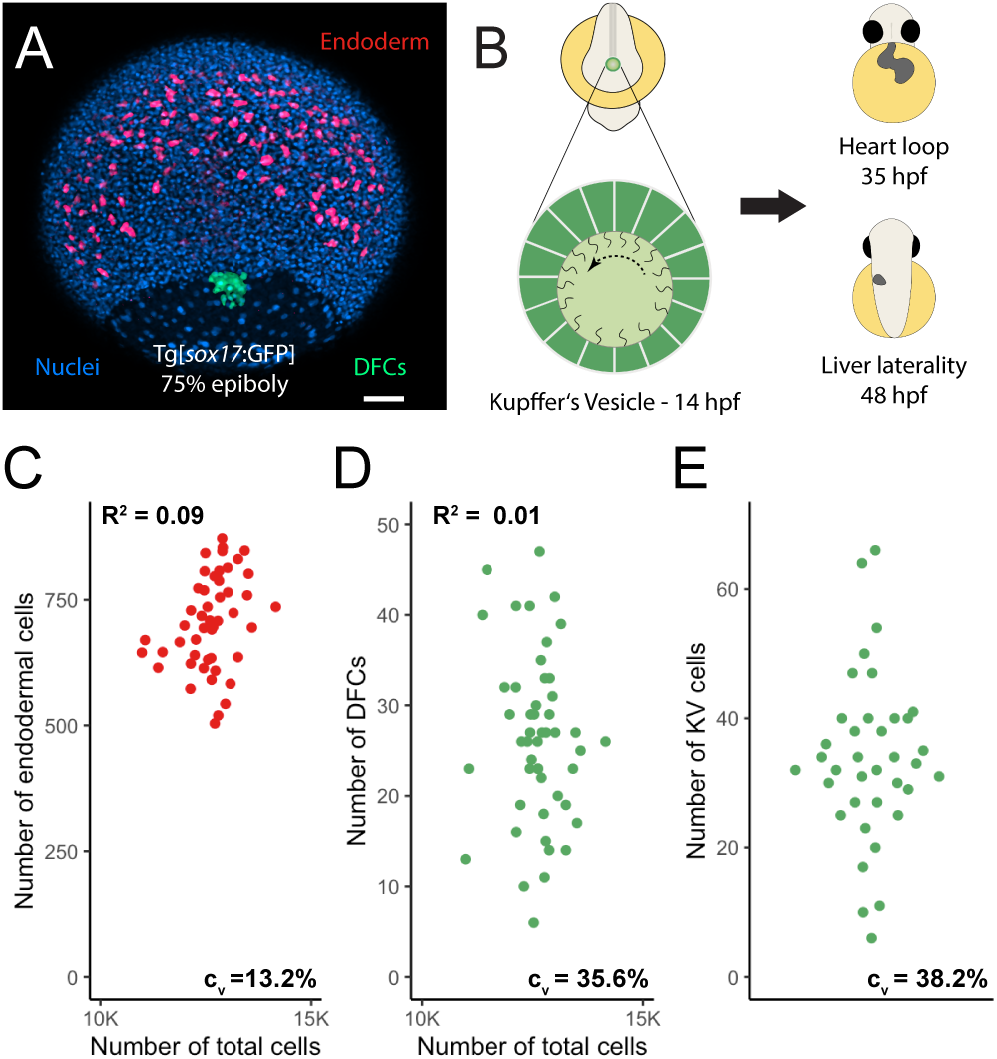
Cell number variability during early embryogenesis. (A) Maximum projection of confocal images of a Tg[*sox17*:GFP] embryo at 75% epiboly stage showing the endoderm (red), DFCs (green) and nuclei (blue). Scale bar = 80μm. (B) Graphical representation of an 8-somite stage embryo and the KV. (C-E) Cell number in endoderm (n = 49), DFCs (n = 49) and KV (n = 37), respectively, for individual embryos. The coefficient of variation (C_V_) is indicated at the bottom right; (C, D) also show the total cell number on the x-axis (C_V_ = 4.8%) and the coefficient of determination (R^2^). Total cell numbers are in good agreement with previous publications (Kobitski et al., 2015).

## Results

To characterize cell number variability during zebrafish early embryogenesis, we counted the number of endodermal cells and DFCs at the 75% epiboly stage using a Tg[*sox17*:GFP] reporter line (Supplemental Figures 1A-D).

We found that the number of endodermal cells exhibits considerable variation, with ~500-800 cells per embryo (Figure 1C). However, the number of DFCs is even more variable, ranging from ~10 to ~50 cells per embryo (Figure 1D). This variation of DFC numbers persists at a later stage when they have formed the KV (Figure 1E, Supplemental Figure 1E), suggesting that no corrective mechanisms are triggered in the animals with particularly high or low numbers of DFCs. These measurements confirm previous publications that reported variable DFC numbers and fluctuations of KV dimensions, albeit at much lower sample numbers (Gokey et al., 2015a, Gokey et al., 2015b). We did not find a positive correlation between total cell number and either the endoderm or the DFCs population (Figures 1C, D), indicating that this variation is not due to staging differences. Furthermore, we found no association between the number of DFCs and the number of endodermal cells (Supplemental Figure 1F), which suggests that the fluctuations of these two cell populations have independent sources.

As an additional technical control, we validated that the GFP cells in the Tg[*sox17*:GFP] line were also expressing the endogenous *sox17* mRNA by detecting both transcripts with fluorescent *in situ* hybridization (Supplemental Figures 1G, H and I). Quantification based on 1-photon microscopy and 2-photon microscopy gave consistent results but, in our setting, the signal to noise ratio was higher in 1-photon microscopy (Supplemental Figure 1J).

The surprisingly large fluctuations in DFC numbers prompted us to investigate possible phenotypic consequences of this variation at later developmental stages. Since the DFCs give rise to the KV, the organ establishing left-right asymmetry, we focused on investigating possible laterality defects. Previous experimental studies, as well as mathematical modeling, suggested that the size of the KV needs to exceed a certain threshold in order to enable reliable left-right patterning (Gokey et al., 2015b; Sampaio et al., 2014). As the heart is the first organ that is formed during zebrafish development, and since its laterality can be assessed easily in live embryos, we examined the percentage of embryos with defective heart laterality (DHL) in different clutches of embryos –Tüpfel long-fin (TL) wild-type strain– at prim-22 stage (Figure 2A; Video 1). We found a DHL average of 3.9% [standard deviation (±): 3.0%, 53 independent observations (IO), total amount of scored embryos (n): 6081]. Even though this is a remarkably high frequency for a wild-type population, this value is in very good agreement with previous reports based on *in situ* hybridization of lower sample numbers ((Wang et al., 2011): 3%, n = 650; (Noël et al., 2013): 6%, n = 387).

**Figure 2.**
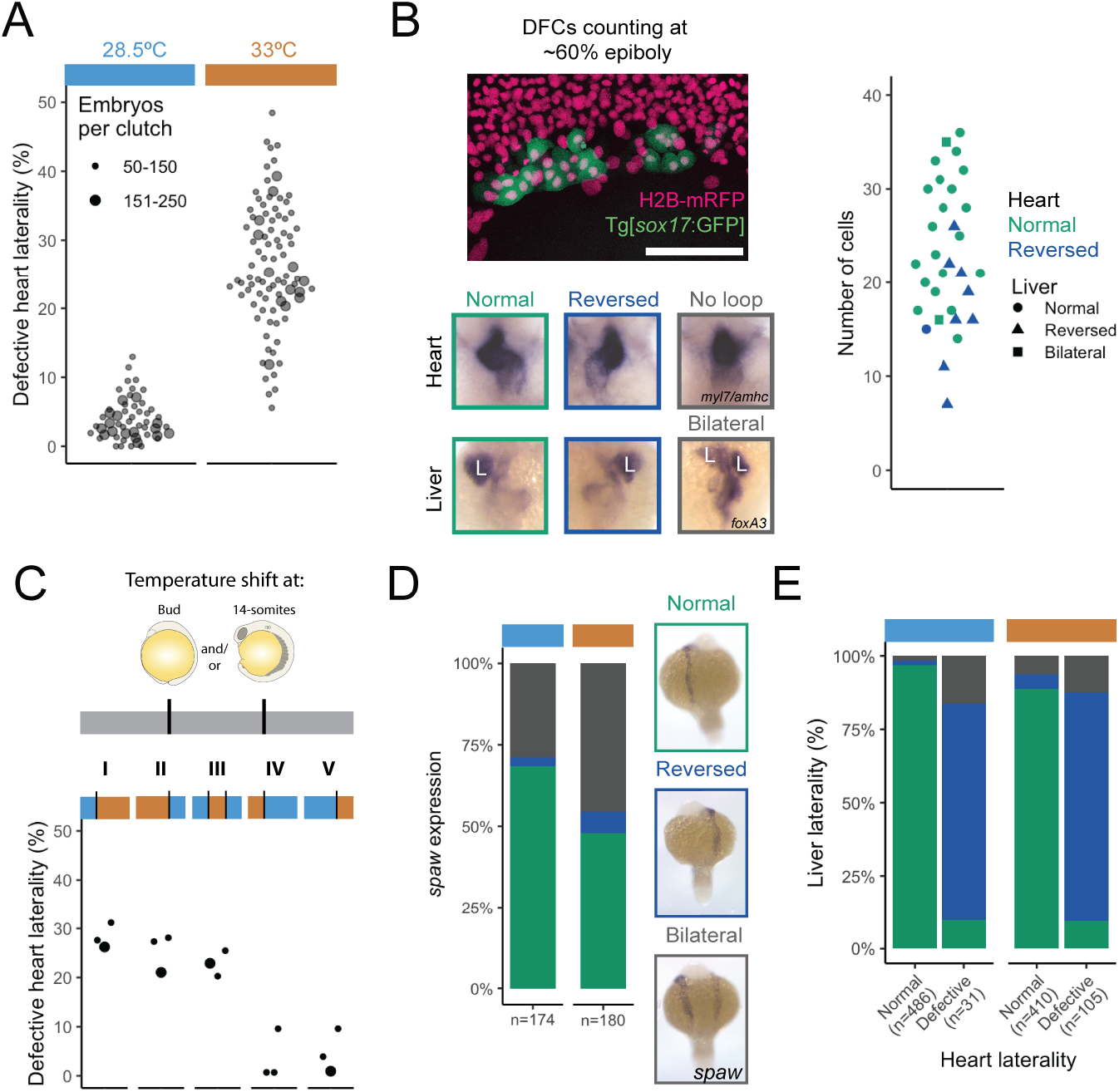
Fluctuations of DFC numbers lead to defects of organ laterality. (A) Defective heart laterality percentages observed at prim-22 stage, including reversed and no heart loop observed in embryos incubated at 28.5°C (blue) and 33°C (orange). The column scatter plots show the percentage of embryos with defective heart laterality per individual clutch analyzed at 28.5°C (marked by the horizontal light blue bar) and 33°C (orange bar). The small circles indicate a clutch size of 50-150 and the bigger ones a range of 151-250 embryos. (B) Confocal z-projection of the dorsal side of a Tg[*sox17*:GFP] (green) live embryo injected with H2B-mRFP (pink). The plot shows the DFC number distribution at 60% epiboly and the resulting heart and liver laterality in the individual embryos (n = 31) assessed by *in situ* hybridization at long pec stage: *myl7* and *amhc* for the heart and *foxA3* for the liver (L). Scale bar = 100μm. (C) Graphical representation of the different temperature shift treatments: the first third of the gray bar represents the period between 1-cell and bud stage, the second until the 14-somite stage, and the third until the period of collection (prim-22 stage). Circle size and incubation temperature as in (A). Treatments: (I) incubation at 28.5°C until Bud Stage, change to 33°C; (II) 33°C until 14-somite, then transferred to 28.5°C; (III) 28.5°C until Bud Stage, shift to 33°C until 14-somite (4.5 hours), then, back to 28.5°C; (IV) 33°C until Bud Stage, change to 28.5°C; (V) 28.5°C until 14-somite, then transferred to 33°C. (D) Relative frequency of different *spaw* expression patterns in embryos incubated at 28.5° and 33°C. *in situ* hybridization photographs of normal (green), reversed (dark blue) and bilateral (gray) *spaw* expression in 18-somite stage embryos are shown to the right. (E) Relative frequency of embryos with normal, reversed or bilateral liver, separated into embryos with normal or defective heart laterality at long pec stage (incubated at 28.5° or 33°C).

We then evaluated whether a change in the environmental conditions could unmask an underlying sensitivity in individuals that would otherwise present normal organ laterality. Previous reports have observed an influence of temperature on the penetrance of zebrafish mutant phenotypes (Imai et al., 2001). The incubation temperature for zebrafish embryos ranges from 25°C to 33°C, with 28.5°C being the standard (Kimmel et al., 1995). Increasing the temperature to 33°C led to a remarkable increase of the DHL average to 26.4% ± 9% (IO = 89, n = 9557) (Figure 2A). We observed no difference in the mortality rate (24.9% ± 13.9% and 24.1% ± 13.8%, at 28.5°C and 33°C, respectively, Supplemental Table 1), and the fraction of abnormal embryos (showing a swollen heart cavity or tail defects, which were excluded from analysis in any condition) remained moderate (4.8% ± 7.5% and 11.7% ± 9.8% at 28.5°C and 33°C, respectively, Supplemental Table 1). Taken together, these observations suggest that establishment of left-right laterality in zebrafish is relatively unstable, which we hypothesized might be due to the variable number of DFCs.

To test this idea, we set out to determine whether the measured fluctuations in DFC numbers are directly linked to the observed laterality variation. To do so, we counted the number of DFCs by live microscopy at 60% epiboly stage and assessed the laterality of the heart and the liver at later stages, after incubation at 33°C. We found that the number of DFCs at 60% epiboly is strongly predictive of laterality defects by long pec stage (Figure 2B), which establishes a direct association between DFC numbers and laterality defects (p-value < 0.02, Fisher’s exact test). Of note, we observed that the fraction of embryos with laterality defects is ~50% for those individuals with less than ~20 DFCs, suggesting that laterality is established randomly (and hence half of the time correctly) in embryos with low DFC number. In summary, we found that an early fluctuation of the size of a small cell population is correlated with a macroscopic defect at later stages, rather than be being corrected over the course of development.

We then aimed to exploit the influence of temperature as a tool to better understand the mechanism of how variability in DFC numbers influences the frequency of heart laterality defects. To this end, we tested several temperature shift treatments spanning early development (1-cell to bud stage), early somitogenesis (bud to 14-somites stage) and organogenesis (following 14-somite stage) (Figure 2C). Interestingly, we found that treating the embryos during early somitogenesis alone, the time at which the KV is formed, was sufficient to generate a similar DHL frequency as with continuous incubation at 33°C (Figure 2C, III). This result suggests that the temperature treatment mostly influences KV function (and hence global leftright patterning). However, temperature does not have a major influence on the number of DFCs (Supplemental Figure 2A), which are determined before somitogenesis, or heart looping, which happens after somitogenesis. To corroborate this hypothesis, we decided to investigate potential defects of global left-right patterning mediated by the KV. Rotational movements by cilia create a directional fluid flow in the extracellular space (Essner et al., 2005). This is thought to activate Nodal signaling in the left lateral plate mesoderm by inducing degradation of the Nodal antagonist *dand5* on the left side and lead to expression of the Nodal ligand *southpaw* (*spaw*) only on the left side of the lateral plate mesoderm (Schneider et al., 2010), which is the molecular signal used to establish organ laterality (Schweickert et al., 2017). Indeed, we found that asymmetric expression of *dand5* was reduced at 33°C compared to 28.5°C (Supplemental Figure 2B, C), suggesting that the elevated temperature interferes with the patterning activity of the KV.

To gain further insight into how the number of cells is related to the function of the KV, we measured the size of the KV as well as the number of cells (Supplemental Figure 2D-F). We found that, as expected, the size of the KV and the number of cells are correlated (R^2^ = 0.78 for 28.5°C, 0.58 for 33°C). However, while the number of cells remained similar, the size of the KV lumen was significantly decreased at 33°C compared to 28.5°C (Supplemental Figure 2E-F). Taken together, these results suggest that an elevated temperature leads to an increase of the cell threshold required for reliable functioning of the KV, which is mediated at least partially via a reduction of the size of the lumen. Of note, we did not observe a change in the probability of heart laterality defects upon a decrease of the incubation temperature to 23°C (Supplemental Figure 2G).

In line with these observations, we found a gradient of expression patterns for the early left-right marker gene *spaw*, ranging from the expected pattern (i.e. *spaw* only on the left) to bilateral expression (*spaw* visible on both sides, although not necessarily at the same level) and complete reversal (i.e. *spaw* only on the right) (Figure 2D). We hypothesized that these defects on the level of *spaw* expression should produce concordant laterality defects in different organs that exhibit left-right asymmetry, such as the heart and liver. Indeed, we found that in most, but not all, cases the embryos with reversed heart laterality also exhibited reversed liver laterality (Figure 2E). We speculate that discordant organ laterality might occur in case of almost perfectly balanced bilateral *spaw* expression (i.e., the same expression levels on both sides), leading to a scenario where in some cases the two organs would develop their laterality largely randomly and independently from each other.

After observing this strong and macroscopically visible phenotype caused by cell number fluctuations at very early developmental stages, we wanted to understand the origin of the DFC number fluctuations. First, we set out to distinguish stochastic fluctuations from genetically determined variation of cell numbers. To identify possible genetic factors, we raised embryos with a reversed heart loop to adulthood and evaluated the DHL frequency in their progeny. As a control, embryos with normal heart laterality were raised as well. To our surprise, we detected no significant differences in either the percentage of reversed heart laterality (Figure 3A) or KV cell number (Supplemental Figure 3A) in the offspring compared to the control group, indicating that this phenotype is not caused by a distinct mutation. While analysis of clones would be required to rule out a genetic component, this result suggests a large stochastic contribution to the percentage of reversed heart laterality and KV cell numbers.

**Figure 3.**
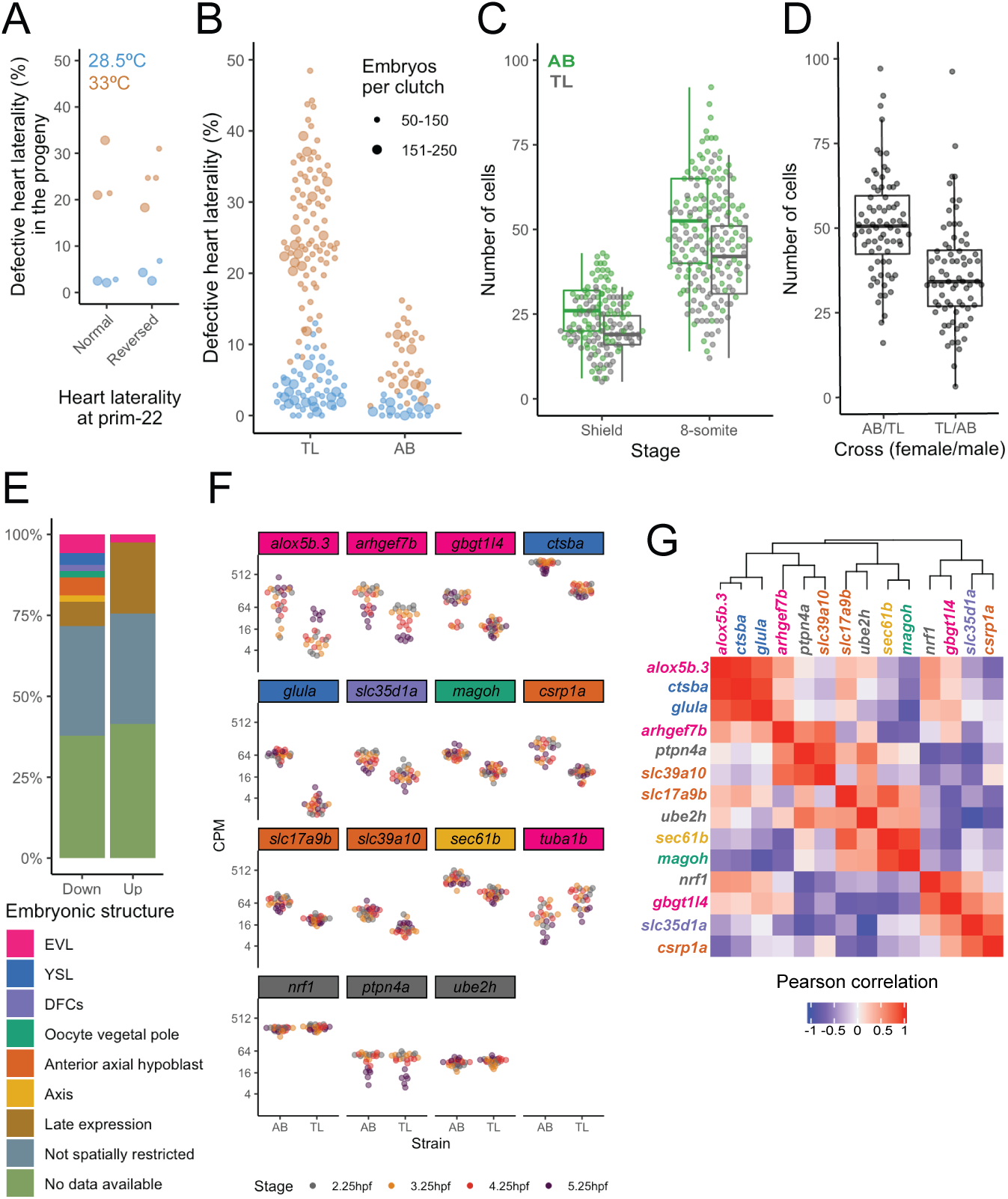
Heart laterality defects are a stochastic phenomenon that is linked to variable deposition of maternal factors. (A) Percentage of embryos with defective heart laterality in the progeny of individuals that showed either normal or reversed heart laterality at prim-22 stage. Incubation temperature 28.5°C (light blue) or 33°C (orange). The small points indicate a clutch size ranging of 50-150 and the bigger points a range of 151-250. (B) Percentage of embryos with defective heart laterality per individual clutch analyzed at 28.5° and 33°C in TL (same data as Figure 2A) and AB embryos. (C) Number of DFCs / KV cells at shield and 8-somite stage; for shield stage: n = 78 and 75 for AB and TL, respectively, p < 0.001; for 8-somite stage: n = 94 and 97 AB and TL, respectively, p < 0.001. (D) KV cell numbers at 8-somite stage in crosses between individual AB and TL males and females, AB/TL versus TL/AB p <0.001 (n=76 for both); AB versus AB/TL p = 0.461; TL versus TL/AB p = 0.01; AB versus TL/AB p <0.001; TL versus AB/TL p < 0.001. The boxplots display the median, the hinges represent the first and third quartiles, the whiskers represent 1.5 of the inter-quartile range from the hinge. (E) Summary of reported embryonic expression patterns for the down- and upregulated genes. (F) Expression levels (in counts per million) for genes that are differentially expressed at all stages. (G) Heatmap showing the pairwise Pearson correlation between the downregulated genes at 2.25hpf for TL embryos. Color scale at the bottom. (F, G) Gene names are color-coded to show the embryonic structure in which they are expressed. Three randomly selected genes with similar expression levels were included as an outgroup (gray).

The seemingly stochastic origin of the observed laterality defects has important conceptual consequences for our interpretation of variable disease phenotypes (see Discussion). However, the stochastic nature of the phenotype numbers makes it more difficult to identify the molecular origin of DFC number fluctuations. We therefore tried to identify general principles that underlie DFC number variability. As both the DFCs and the surrounding dorsal domain are induced by high levels of Nodal in a non-cell-autonomous manner (Hagos and Dougan, 2007; Oteíza et al., 2008), we reasoned that there could be a direct correlation between the number of cells forming the dorsal organizer and the DFCs number; however, we didn’t find such an association (Supplemental Figures 3B, C). Furthermore, we only found a weak association between maternal age and DHL frequency at 28.5°C (Supplemental Figure 3D).

Inspired by reports that described DHL as a spontaneous strain-dependent defect (Malicki et al., 2011), as well as previous observations of strain-dependent differences in KV dimensions (Gokey et al., 2015b), we next compared the frequencies of reversed heart laterality between embryos with TL genetic background (the one used so far) versus AB. Interestingly, we found significantly lower DHL frequencies for AB embryos at 28.5°C (1.3% ± 1.3%, IO = 22, n = 2714) and 33°C (7.7% ± 4.1%, IO = 34, n = 3353) compared to TL (Figure 3B, Supplemental Table 1). In line with this observation, we found that the number of DFCs and KV cells was higher in AB versus TL (Figure 3C), and that the antero-posterior cilia distribution (AP ratio) is more asymmetric in AB than in TL (Supplemental Figures 3E, F).

The strain-specific differences in DFC numbers gave us a handle to explore the molecular mechanism in more detail. Specifically, we hypothesized that the fluctuations might be maternally controlled, since DFCs begin to be emerge 1 hour after maternal-to-zygotic transition, at 4 hpf (Oteíza et al. 2008). To test this hypothesis, we crossed AB females with TL males and vice versa. Indeed, we found that the number of KV cells depends mostly on the mother’s genetic background, as the cell numbers of the two hybrid crosses are different from each other [the AB (female) / TL (male) cross resembles the AB/AB cross, and the TL (female) / AB (male) cross is more similar to the TL/TL cross, although the latter two distributions still differ with a p-value < 0.01] (Figure 3C, D). Consequently, a major source of the observed fluctuations in the embryo must lie in the development of the oocytes in the mother.

When comparing TL and AB embryos by RNA-seq, we found 94 genes that were consistently differentially expressed in the period before and after the zygotic genome activation (2.25 hpf and 3.25 hpf, respectively) as well as the time window during which the DFCs are specified (4.25-5.25 hpf) (Figure 3E, Supplemental Figure 4B, Supplemental Tables 2 and 3). 12 of the differentially expressed genes are reported to have an early expression in specific cell types or structures (Figures 3E, F, Supplemental Figures 4C, D, F and H). It is very likely that most of these differentially expressed genes are related to processes other than DFC specification that differ between the two strains. However, we were intrigued to find that 7 of these 12 genes with a reported early spatial pattern are expressed either the DFCs, in cell types related to their specification – the enveloping layer (EVL), from which the DFCs are derived (Cooper and D’Amico, 1996; Hagos and Dougan, 2007; Oteíza et al., 2008) – and the yolk syncytial layer (YSL), which remains connected to the DFCs until 4hpf (Amack and Yost, 2004; Cooper and D’Amico, 1996). Of note, embryo-to-embryo variability of transcript levels for these candidate genes is lower than the expression difference between the two strains (Figure 3E), suggesting that the observed phenomenon cannot be explained by a simple thresholding mechanism on the level of transcript abundance.

We found a high correlation in the expression levels of several of the differentially expressed YSL/EVL genes across the TL embryos (Figure 3G, Supplemental Figures 4E, G and I). This raises the possibility that variation of some upstream factor (or a combination of factors) acting during oocyte development may be responsible for the observed variable but correlated expression. Interestingly, the mutant for one of these genes, *ctsba*, has been reported to have a reduced number of EVL cells (Langdon et al., 2016). However, it is unlikely that the genes identified here are the only ones responsible for variable DFC numbers, since additional factors (e.g. fluctuations in protein levels, or structural differences in the oocyte) may contribute equally. Furthermore, additional genes without a reported YSL/EVL expression might also be involved in this phenomenon. In summary, these experiments suggest that maternal effects, possibly linked to deposition of factors involved in YSL/EVL specification and function, may be responsible for the observed fluctuations in DFC numbers. However, additional experiments would be required to functionally link transcript levels of specific genes to the observed phenotype.

## Discussion

We found that fluctuations of a small cell population generated during the earliest stages of embryogenesis (the DFCs) are not corrected over the course of development, but instead persist and give rise to a macroscopic phenotype (defects of organ laterality) if the embryos are incubated at 33°C. While it was reported before that left-right patterning in wild-type zebrafish is surprisingly variable (Noël et al., 2013; Wang et al., 2011) and that there is a size threshold for reliable functioning of the KV (Gokey et al., 2015b; Sampaio et al., 2014), we now show that these phenomena are linked and originate in fluctuations of DFC numbers that are at least partially caused by variable maternal factors.

Our data suggests that both genetic and stochastic components are involved in DFC variability: If we compare animals from different strains, genetic factors are the dominant effect, but if embryos of a single strain are analyzed, a seemingly stochastic effect becomes dominant. Analysis of clones would be required to truly quantify the degree of stochastic fluctuations. However, our findings indicate that this phenomenon is not caused by a distinct mutation.

Here we show that the number of cells in the KV is important for proper function. However, other studies have shown that the internal architecture is equally critical, since a higher density of cilia in the anterio-dorsal region is needed to create a proper leftward flow (Kreiling et al., 2007; Okabe et al., 2008). We did observe differences in the AP ratio between AB and TL strains but no correlation to the cilia number. If and how variation in DFC number is linked to morphological asymmetries in the KV is an interesting question, which would require additional live experiments.

The sizes of progenitor populations in vertebrate embryos are inherently variable due to environmental, genetic and stochastic fluctuations. Without feedback control, the size of the final cell population would dependent linearly on the size of the progenitor pool and hence be highly volatile (Lander et al., 2009). It is therefore remarkable that in this case there seem to be no corrective mechanisms that reduce cell number variability and thereby ensure robust leftright patterning. Furthermore, while mouse embryos pass a size checkpoint at around the time of gastrulation (Lewis and Rossant, 1982; Snow and Tam, 1979), we speculate that the externally developing zebrafish embryos may not have to pass such a checkpoint and might instead have evolved to maximize the speed of development, even at the cost of occasional laterality defects.

We investigated the consequences of developmental variability in a specific cell population, the DFCs, and it remains to be answered how general such phenomena are in other cell populations. Interestingly, primordial germ cells numbers are also variable and higher in AB versus TL embryos (Weidinger et al., 1999). How variable vertebrate development is, and which degree of fluctuations is still compatible with normal development, are important open questions, which will require novel experimental approaches such as high-throughput lineage tracing (Alemany et al., 2018; Raj et al., 2018; Spanjaard et al., 2018).

While stochastic developmental defects of this frequency and severity are probably not very common in wild-type mammals, it is likely that similar mechanisms as the one described here may underlie incomplete penetrance of mutations in model organisms (Burga et al., 2011; Raj et al., 2010) as well as in humans. Furthermore, there is increasing evidence that disease phenotypes that manifest late in life, like Alzheimer’s disease, may already originate in development (Arendt et al., 2017). A similar phenomenon may occur in type I diabetes – an emerging view is that the initial pool of beta cells at the end of development is variable between individuals, leading to increased disease susceptibility in a subset of the population (Atkinson et al., 2015).We therefore speculate that early stochastic fluctuations of progenitor cell pools might at least partially contribute to a variety of human disease phenotypes.

## Acknowledgements

We acknowledge support by the MDC/BIMSB core facilities for zebrafish and advanced light microscopy (especially Anca Margineanu), Jana Richter for help with animal experiments and Nancy Coconi-Linares for artwork. Work in J.P.J.’s laboratory was funded by a European Research Council Starting Grant (ERC-StG 715361 SPACEVAR). R.M.A. was supported by a CONACyT (Mexico) postdoctoral fellowship (CVU 269440).

## Author contributions

J.P.J. and R.M.A. conceived and designed the project. R.M.A performed experiments and analyzed the data, with support by P.O.C. and R.S.. J.P.J. guided experiments and data analysis. All authors discussed and interpreted the results. R.M.A. and J.P.J. wrote the manuscript, with input from P.O.C..

## Declaration of interests

The authors declare no competing interests.

## Methods

### Zebrafish Husbandry and Staging

Adult zebrafish were maintained and bred under standard conditions. Embryos were left to develop in egg water (0.6 g/L dissolved in dH2O; Red Sea Salt, Red Sea, containing methylene blue, 0.002 g/L) to the desired stage at 28.5°C unless otherwise stated. Staging was done based on (Kimmel et al., 1995). Zebrafish were bred, raised, maintained, and handled in accordance with the guidelines of the Max-Delbrück Center for Molecular Medicine and the local authority for animal protection (Landesamt für Gesundheit und Soziales, Berlin, Germany) for the use of laboratory animals based on the current version of German law on the protection of Animals and EU directive 2010/63/EU on the protection of animals used for scientific purposes.

### Zebrafish strains

AB and Tüpfel Long Fin (TL) strains were obtained from the European Zebrafish Research Center (EZRC, 1175 and 1174, respectively). The TL strain was used as a default, unless stated otherwise. The Tg[*sox17*:GFP] strain was reported in (Sakaguchi et al., 2006). To change its genetic background to TL, the fish were crossed with TL fish. By crossing the resulting males with TL females, we obtained similar DHL percentages as in the TL wild-type strain.

### Immunolocalization

Embryos at the desired stage were fixed overnight at 4°C in PFA 4% in PBS. The following day, they were washed 3x with PBSTx (1x PBS with TritonX 1%) for 5 minutes, dechorionated and incubated in blocking buffer (2% BSA, 5% Goat Serum in PBSTx) for 2 hours at room temperature (RT), followed by an overnight incubation at 4°C with one or a combination of the following primary antibodies diluted in blocking buffer: mouse anti-acetylated Tubulin (1:200, Sigma Aldrich T6793), chicken and rabbit anti-GFP (1:1000, Abcam ab13970 and ab290, respectively), rabbit anti-Flh (1:200, Sigma Aldrich SAB2702443) and rabbit anti-Laminin-α1β1γ1 (1:100, Sigma Aldrich L9393). On the next day, they were washed 3x with PBSTx and 1x with blocking buffer, 30 minutes each at RT, followed by an overnight incubation at 4°C with Hoechst (1:1000, Thermo Fisher 62249) and secondary antibody diluted in blocking buffer (1:200): goat anti-chicken Alexa-488 (Thermo Fisher A11039), goat anti-mouse Alexa-647 (Thermo Fisher A32728) or goat anti-rabbit Alexa-568 (Thermo Fisher A11011). Finally, 3x PBSTx washes, 30 minutes each at RT.

### Imaging

Endodermal cells, DFCs and total cell number in 75% epiboly embryos: after anti-GFP immunolocalization, the embryos were washed with 50% glycerol/50% PBSTx for ten minutes and then with 100% glycerol overnight at 4°C. For flat mounting, a layer of laboratory labeling tape (13mm wide) was attached to each side of a coverslip to make a bridge and leave enough room for the flat embryo (~120μm). A single embryo was put on a coverslip with a drop of glycerol. For flattening, a closed forceps was introduced in the vegetal pole and then opened to break and split the yolk cell in half. As many yolk granules as possible were removed to avoid damaging the blastoderm, then another coverslip was put on top. The tissue was oriented facing the bottom to be imaged with an SP8 inverted confocal microscope. After obtaining an image stack for the endodermal and total cell number, another stack was obtained for the DFC region with lower laser power and higher zoom factor, since the GFP signal intensity in these cells is considerably higher than in the endodermal cells. SP8 microscope acquisition parameters: 20x multi-immersion objective, format = 1024 x 1024, speed = 600, zoom factor = 0.75 (except for DFCs, 2 or 4 when all cells fit), line average = 2, Z-step size = 3, gating = 0.3-6 for both channels, laser intensity Z-compensation for each image, tiling = 2×2 in most images, with 15% overlapping. The images were automatically stitched with the Leica software.

For imaging DFCs at shield, cilia at 8-somite stage and total amount of cells at different stages (2.25-5.25hpf), the embryos were fixed inside the chorion, mounted in 1% low melting point agarose, and imaged on an upright Zeiss 880 confocal microscope, with the following settings: 20x water-dipping objective, format = 512 x 512, speed = 8, zoom factor = 1, line average = 2, Z-step size = 3. For counting the total number of cells (as well as for a comparison of DFC quantification to 1-photon microscopy), a 2-photon Chameleon laser was used; for the total cell count, tiling = 2×2, with 15% overlapping was done and the images were automatically stitched with the Zeiss software.

### Live imaging

For KV measurements at 8-somite stage, embryos incubated at 28.5°C or 33°C were mounted on a 1.5% agarose injection dish with little liquid. The embryos were oriented inside the chorion to image the KV. Afterwards, 3% methylcellulose in E3 embryo medium (5mM NaCl, 0.17mM KCl, 0.33mM CaCl2, 0.33mM MgSO4) was added to cover the embryos and, thus, retain the order of the individual embryos. The embryos were incubated at the initial temperature until prim-22 for heart laterality analysis. Per session, around 30 embryos were imaged for each condition.

For DFC counting, individual Tg[*sox17*:GFP] (injected with ~60 pg of H2B-mRFP mRFP1 at one-cell stage) shield stage embryos were transferred to glass-bottom Petri dishes, most of the liquid was removed, and ~0.5 mL of 3% methylcellulose in E3 embryo medium was added. Then, the embryos were manually dechorionated with forceps and oriented with the shield facing the bottom of the dish for imaging on a Leica SP8 confocal microscope. After imaging, the dish was filled with E3 embryo medium and incubated at 33°C until long pec stage and fixed for *in situ* hybridization.

### *in situ* hybridization

Whole-mount in situ hybridizations were performed essentially as described previously (Thisse and Thisse, 2008). The following probes were used: *foxA3* and *myl7* (Noël et al., 2013); and *spaw* and *dand5* (Sampaio et al., 2014).The *amhc* probe was kindly provided by Daniela Panáková. For fluorescent *in situ* hybridization, the RNAscope probes Dr-sox17-C3 and EGFP (Advanced Cell Diagnostics 400281 and 494711-C3, respectively) were used and the detection was done as described previously (Gross-Thebing et al., 2014) with the RNAscope Fluorescent Multiplex Reagent Kit (Advanced Cell Diagnostics 320850) and imaged on a confocal microscope.

### mRNA synthesis

The H2B-mRFP1 mRNA was synthesized using the mMESSAGE mMACHINE kit (Thermo Fisher AM1345) according to the manufacturer’s recommendations.

### Temperature shift treatment

After collection, the embryos were put immediately in warmed water, either to 28.5°C or 33°C. For the temperature shift, the embryos were transferred to a Petri dish filled with water at the desired temperature.

### Single embryo RNA-seq

A total of 6 AB and 6 TL embryos were individually collected per stage (2.25, 3.25, 4.25 and 5.25 hpf) in LoBind tubes (Eppendorf) in two independent experiments (each of which contained half of the samples for each condition). 0.5mL Trizol and 0.5μl GlycoBlue were added to each samples, and RNA extraction was carried according to the manufacturer’s recommendations. Each sample was barcoded, pooled and the libraries were prepared according to the CEL-seq2 protocol (Hashimshony et al., 2016) with different RPI indices for each timepoint and paired-end sequenced on an Illumina NextSeq 500 (read length 12 nt for barcode read and 63 nt for transcript read).

### Sequencing data analysis

Basecalling was done with bcl2fastq v2.19.0.316. The resulting reads were demultiplexed with *scruff* (Wang et al., 2019). Mapping was done with STAR 2.7.1a (Dobin et al., 2013) using quantMode GeneCounts and the DanRer11v96 transcriptome as reference. Differentially expressed genes were obtained with edgeR (Robinson et al., 2009). Genes with a False Discovery Rate (FDR) < 0.01, log2 Counts Per Million > 4 and log2 Fold Change < −1 for downregulated and > 1 for upregulated genes were considered differentially expressed. Spatial and temporal expression was obtained on The Zebrafish Information Network (Bradford et al., 2011).

### Pairwise correlation for gene levels

To avoid a bias due to outliers, we applied a logarithmic transformation of the counts per million values for each sample (6 per strain) for the desired downregulated genes in TL embryos; these values were used to obtain a matrix of Pearson correlations for gene pairs and to generate the heatmaps.

### Plots

All plots were generated in R using the following additional libraries: ggplot2 graphing package, ggbeeswarm for column scatter plots, ggExtra for marginal histograms and ComplexHeatMap for heatmaps. Volcano plots were generated with VolcaNoseR (Goedhart and Luijsterburg, 2020).

### Estimation of KV cell number thresholds

We performed the following back-of-the-envelope calculation to determine whether the KV cell number threshold is the same in AB and TL animals: In Figure 2B, we found that a threshold value of ~20 DFCs is required for reliable left/right patterning at 33°C. The DFCs divide on average about once until they form the KV (Figure 1D,E), which would lead to a threshold of ~40 KV cells. A KV cell number threshold of around 30 had been postulated previously at normal temperature (Sampaio et al., 2014). It makes sense that our threshold is a little higher, since we observed an increased sensitivity at elevated temperature (Figure 2A). Using the data from Figure 3C at the 8 somite stage, we find 20.6% of AB embryos and 37.1% of TL embryos have KV cell numbers below 40. Assuming a 50/50 chance of correct versus inverted organ laterality in the absence of reliable left/right signaling, we would hence expect 10.3% of inverted hearts in AB, and 18.6% in TL. These numbers are within the range of observed values in Figure 3B (7.7% ± 4.1% for AB and 26.4% ± 9% for TL). However, it’s important to point out that other factors might also contribute to the observed discrepancy between AB and TL, such as differences in the expansion of the KV lumen, or differences in cilia density in sub-regions of the KV.

### Cell counting

The endodermal cells were counted with the Imaris (Bitplane) Surface module: background subtraction: on, diameter of largest sphere: 15 μm, splicing touching: on, seed points diameter: 8 μm. For each sample, the number was visually corrected. The total cell number was estimated with the Fiji (Schindelin et al., 2012) ITCN plugin. A Z-projection was obtained for each stack, the ITCN parameters were: width 12, minimum distance 6, threshold: 0.5.

For quantification of dorsal domain cells and DFCs, cell number was estimated manually for cilia with the Cell Counter Fiji plugin. The cilia number was used as a proxy for the KV cell number, since the cells are monociliated. DFCs numbers were first estimated with ITCN (width: 18, minimum distance: 9, threshold: 0.5) and visually corrected afterwards. For the estimation of the total number of cells, the cells were counted with Imaris, using the same parameters as described above.

The Flh positive cells were counted with the ITCN plugin (same parameters as described above for the DFCs). In all cases the embryos were imaged immediately after immunolocalization. The number of dorsal domain cells was obtained by subtracting the DFC number from the number of Flh positive cells, since Flh is also expressed in the DFCs.

### Anteroposterior ratio

Using Laminin-α1β1γ1λ localization as a reference for the anterior position of the KV, we counted the KV cilia in the anterior and posterior halves. The AP ratio was calculated as the ratio between these values.

### Statistics

The coefficients of determination were obtained with R. The p-values were obtained with a randomization test done with PlotsOfDifferences (Goedhart, 2019). All statistical parameters, including sample numbers and median are shown in the figures and described in the figure legends or in the main text.

## Data Availability

The RNA-seq data generated in this study is available on GEO, accession code GSE153621, https://www.ncbi.nlm.nih.gov/geo/query/acc.cgi?acc=GSE153621.

## Supplemental items

**Video 1. Heart loop phenotypes observed at prim-22 stage.** Heart laterality phenotypes on live embryos; normal/dextral loop (d-loop); defective includes: reversed/sinistral loop (s-loop) and no loop, shown side by side on a ventral view.

**Supplemental Table 1. Defective heart laterality numbers per observation.** Genetic background, strain, mother age, temperature of incubation; number of dead, normal, reversed, no discernible heart loop and with abnormal morphology; total amount scored, alive and with normal morphology; percentage of dead, defective heart laterality (reversed + no loop), reversed heart laterality (RHL) and abnormal. The age of the mother was not determined for some observations (ND), hence, these experiments were not used for the correlation DHL% versus mother age.

**Supplemental Table 2. Shared differentially expressed genes.** Gene name, Ensembl ID, function, earliest expression, spatial expression, differential expression and embryonic structure information from ZFIN; log2FC, log2CPM, p-value and FDR for each stage selected for strain comparison.

**Supplemental Table 3. Normalized counts for DE genes.** Counts per million (CPM) levels for the shared differentially expressed and the outgroup genes for all stages.

## Notes

https://www.ncbi.nlm.nih.gov/geo/query/acc.cgi?acc=GSE153621

### Competing Interest Statement

The authors have declared no competing interest.

## References

Alemany, A., Florescu, M., Baron, C.S., Peterson-Maduro, J., and Van Oudenaarden, A. (2018). Whole-organism clone tracing using single-cell sequencing. Nature 556, 108–112.

Alexander, J., Rothenberg, M., Henry, G.L., and Stainier, D.Y. (1999). Casanova plays an early and essential role in endoderm formation in zebrafish. Developmental Biology 215, 343–357.

Amack, J.D., and Yost, HJ. (2004). The T Box Transcription Factor No Tail in Ciliated Cells Controls Zebrafish Left-Right Asymmetry. Current Biology 128, 189–190.

Arendt, T., Stieler, J., and Ueberham, U. (2017). Is sporadic Alzheimer’s disease a developmental disorder? Journal of Neurochemistry 143, 396–408.

Atkinson, M., von Herrath, M., Powers, A.C., and Clare-Salzler, M. (2015). Current Concepts on the Pathogenesis of Type 1 Diabetes d Considerations for Attempts to Prevent and Reverse the Disease. Diabetes Care 38, 979–988.

Barkai, N., and Shilo, B.Z. (2009). Robust generation and decoding of morphogen gradients. Cold Spring Harbor Perspectives in Biology 1, 1–15.

Bradford, Y., Conlin, T., Dunn, N., Fashena, D., Frazer, K., Howe, D.G., Knight, J., Mani, P., Martin, R., Moxon, S.A.T., et al. (2011). ZFIN: Enhancements and updates to the zebrafish model organism database. Nucleic Acids Research 39, 822–829.

Briscoe, J., and Small, S. (2015). Morphogen rules: Design principles of gradient-mediated embryo patterning. Development 142, 3996–4009.

Burga, A., Casanueva, M.O., and Lehner, B. (2011). Predicting mutation outcome from early stochastic variation in genetic interaction partners. Nature 480, 250–253.

Cooper, M.S., and D’Amico, L. a (1996). A cluster of noninvoluting endocytic cells at the margin of the zebrafish blastoderm marks the site of embryonic shield formation. Developmental Biology 180, 184–198.

Dobin, A., Davis, C.A., Schlesinger, F., Drenkow, J., Zaleski, C., Jha, S., Batut, P., Chaisson, M., and Gingeras, T.R. (2013). STAR: Ultrafast universal RNA-seq aligner. Bioinformatics 29, 15–21.

El-Brolosy, M.A., Kontarakis, Z., Rossi, A., Kuenne, C., Günther, S., Fukuda, N., Kikhi, K., Boezio, G.L.M., Takacs, C.M., Lai, S.L., et al. (2019). Genetic compensation triggered by mutant mRNA degradation. Nature 568, 193–197.

Essner, J.J., Amack, J.D., Nyholm, M.K., Harris, E.B., and Yost, H.J. (2005). Kupffer’s vesicle is a ciliated organ of asymmetry in the zebrafish embryo that initiates left-right development of the brain, heart and gut. Development 132, 1247–1260.

Gaspar, P., Arif, S., Sumner-rooney, L., Kittelmann, M., Bodey, A.J., Stern, D.L., Nunes, M.D.S., and Mcgregor, A.P. (2020). Characterization of the Genetic Architecture Underlying Eye Size Variation Within Drosophila melanogaster and Drosophila simulans. G3 10, 1005–1018.

Goedhart, J. (2019). PlotsOfDifferences - a web app for the quantitative comparison of unpaired data. BioRxiv.

Goedhart, J., and Luijsterburg, M.S. (2020). VolcaNoseR is a web app for creating, exploring, labeling and sharing volcano plots. Scientific reports 10:20560.

Gokey, J.J., Dasgupta, A., and Amack, J.D. (2015a). The V-ATPase accessory protein Atp6ap1b mediates dorsal forerunner cell proliferation and left-right asymmetry in zebrafish. Developmental Biology 407, 115–130.

Gokey, J.J., Ji, Y., Tay, H.G., Litts, B., and Amack, J.D. (2015b). Kupffer’s vesicle size threshold for robust left-right patterning of the zebrafish embryo. Developmental Dynamics 22–33.

Gross-Thebing, T., Paksa, A., and Raz, E. (2014). Simultaneous high-resolution detection of multiple transcripts combined with localization of proteins in whole-mount embryos. BMC Biol. 12, 55.

Hagos, E.G., and Dougan, S.T. (2007). Time-dependent patterning of the mesoderm and endoderm by Nodal signals in zebrafish. BMC Developmental Biology 18, 1–18.

Hashimshony, T., Senderovich, N., Avital, G., Klochendler, A., de Leeuw, Y., Anavy, L., Gennert, D., Li, S., Livak, K.J., Rozenblatt-Rosen, O., et al. (2016). CEL-Seq2: Sensitive highly-multiplexed single-cell RNA-Seq. Genome Biology 17, 1–7.

Hunt, E.R., Dornan, C., Sendova-franks, A.B., and Franks, N.R. (2018). Asymmetric ommatidia count and behavioural lateralization in the ant Temnothorax albipennis. Scientific Reports 1–11.

Imai, Y., Gates, M. a, Melby, a E., Kimelman, D., Schier, a F., and Talbot, W.S. (2001). The homeobox genes vox and vent are redundant repressors of dorsal fates in zebrafish. Development 128, 2407–2420.

Kimmel, C.B., Ballard, W., Kimmel, S.R., Ullman, B., and Schilling, T.F. (1995). Stages of Embryonic Development of the Zebrafish. Developmental Dynamics 203, 255–310.

Kobitski, A.Y., Otte, J.C., Takamiya, M., Schäfer, B., Mertes, J., Stegmaier, J., Rastegar, S., Rindone, F., Hartmann, V., Stotzka, R., et al. (2015). An ensemble-averaged, cell densitybased digital model of zebrafish embryo development derived from light-sheet microscopy data with single-cell resolution. Scientific Reports 5, 1–10.

Kreiling, J.A., Prabhat, Williams, G., and Creton, R. (2007). Analysis of Kupffer’s vesicle in zebrafish embryos using a cave automated virtual environment. Developmental Dynamics 236, 1963–1969.

Lander, A.D., Gokoffski, K.K., Wan, F.Y.M., Nie, Q., and Calof, A.L. (2009). Cell Lineages and the Logic of Proliferative Control. 7.

Langdon, Y.G., Fuentes, R., Zhang, H., Abrams, E.W., Marlow, F.L., and Mullins, M.C. (2016). Split top: a maternal cathepsin B that regulates dorsoventral patterning and morphogenesis. Development 143, 1016–1028.

Lewis, N.E., and Rossant, J. (1982). Mechanism of size regulation in mouse embryo aggregates. Journal of Embryology and Experimental Morphology Vol. 72, 169–181.

Malicki, J., Avanesov, A., Li, J., Yuan, S., and Sun, Z. (2011). Analysis of Cilia Structure and Function in Zebrafish. Methods in Cell Biology, 101, 39–74.

Megason, Sean G. 2009. In Toto Imaging of Embryogenesis with Confocal Time-Lapse Microscopy. Mol Biol. Vol. 546.

Melby, A.E., Warga, R.M., and Kimmel, C.B. (1996). Specification of cell fates at the dorsal margin of the zebrafish gastrula. Development 122, 2225–2237.

Noël, E.S., Verhoeven, M., Lagendijk, A.K., Tessadori, F., Smith, K., Choorapoikayil, S., den Hertog, J., and Bakkers, J. (2013). A Nodal-independent and tissue-intrinsic mechanism controls heart-looping chirality. Nature Communications 4, 2754.

Okabe, N., Xu, B., and Burdine, R.D. (2008). Fluid dynamics in zebrafish Kupffer’s vesicle. Developmental Dynamics 237, 3602–3612.

Oteíza, P., Köppen, M., Concha, M.L., and Heisenberg, C.-P. (2008). Origin and shaping of the laterality organ in zebrafish. Development 135, 2807–2813.

Raj, A., Rifkin, S. a, Andersen, E., and van Oudenaarden, A. (2010). Variability in gene expression underlies incomplete penetrance. Nature 463, 913–918.

Raj, B., Wagner, D.E., McKenna, A., Pandey, S., Klein, A.M., Shendure, J., Gagnon, J.A., and Schier, A.F. (2018). Simultaneous single-cell profiling of lineages and cell types in the vertebrate brain. Nature Biotechnology 36, 442–450.

Ramaekers, A., Claeys, A., Kapun, M., Mouchel-Vielh, E., Potier, D., Weinberger, S., Grillenzoni, N., Dardalhon-Cuménal, D., Yan, J., Wolf, R., et al. (2019). Altering the Temporal Regulation of One Transcription Factor Drives Evolutionary Trade-Offs between Head Sensory Organs Article Altering the Temporal Regulation of One Transcription Factor Drives Evolutionary Trade-Offs between Head Sensory Organs. Developmental Cell 50, 780–792.

Robinson, M.D., McCarthy, D.J., and Smyth, G.K. (2009). edgeR: A Bioconductor package for differential expression analysis of digital gene expression data. Bioinformatics 26, 139–140.

Sakaguchi, T., Kikuchi, Y., Kuroiwa, A., Takeda, H., and Stainier, D.Y.R. (2006). The yolk syncytial layer regulates myocardial migration by influencing extracellular matrix assembly in zebrafish. Development 133, 4063–4072.

Sampaio, P., Ferreira, R.R., Guerrero, A., Pintado, P., Tavares, B., Amaro, J., Smith, A.A., Montenegro-Johnson, T., Smith, D.J., and Lopes, S.S. (2014). Left-right organizer flow dynamics: How much cilia activity reliably yields laterality? Developmental Cell 29, 716–728.

Schindelin, J., Arganda-Carreras, I., Frise, E., Kaynig, V., Longair, M., Pietzsch, T., Preibisch, S., Rueden, C., Saalfeld, S., Schmid, B., et al. (2012). Fiji: an open-source platform for biological-image analysis. Nature Methods 9, 676–682.

Schneider, I., Schneider, P.N., Derry, S.W., Lin, S., Barton, L.J., Westfall, T., and Slusarski, D.C. (2010). Zebrafish Nkd1 promotes Dvl degradation and is required for left-right patterning. Developmental Biology 348, 22–33.

Schweickert, A, Ott, T, Kurz, S, Tingler, M, M, M., Fuhl, F, and Blum, M (2017). Vertebrate Left-Right Asymmetry: What Can Nodal Cascade Gene Expression Patterns Tell Us? JCDD 5, 1.

Shah, G., Thierbach, K., Schmid, B., Waschke, J., Reade, A., Hlawitschka, M., Roeder, I., Scherf, N., and Huisken, J. (2019). Multi-scale imaging and analysis identify pan-embryo cell dynamics of germlayer formation in zebrafish. Nature Communications 10, 5753.

Snow, M.H.L., and Tam, P.P.L. (1979). Is compensatory growth a complicating factor in mouse teratology? Nature 279, 555–557.

Spanjaard, B., Hu, B., Mitic, N., Olivares-Chauvet, P., Janjuha, S., Ninov, N., and Junker, J.P. (2018). Simultaneous lineage tracing and cell-type identification using CRISPR-Cas9-induced genetic scars. Nature Biotechnology 2018.

Thisse, C., and Thisse, B. (2008). High-resolution in situ hybridization to whole-mount zebrafish embryos. Nature Protocols 3, 59–69.

Wang, G., Cadwallader, A.B., Jang, D.S., Tsang, M., Yost, H.J., and Amack, J.D. (2011a). The Rho kinase Rock2b establishes anteroposterior asymmetry of the ciliated Kupffer’s vesicle in zebrafish. Development 138, 45–54.

Wang, Z., Hu, J., Johnson, W.E., and Campbell, J.D. (2019). Scruff: An R/Bioconductor package for preprocessing single-cell RNA-sequencing data. BMC Bioinformatics 20, 1–9.

Warga, R.M., and Kane, D.A. (2018). Wilson cell origin for kupffer’s vesicle in the zebrafish. Developmental Dynamics 247, 1057–1069.

Warga, R.M., and Nüsslein-Volhard, C. (1999). Origin and development of the zebrafish endoderm. Development 126, 827–838.

Weidinger, Gilbert, Uta Wolke, Marion Köprunner, Michael Klinger, and Erez Raz. 1999. “Identification of Tissues and Patterning Events Required for Distinct Steps in Early Migration of Zebrafish Primordial Germ Cells.” Development 126 (23): 5295–5307.

