## Supplemental Figures for "Variability of an early developmental cell population underlies stochastic laterality defects"

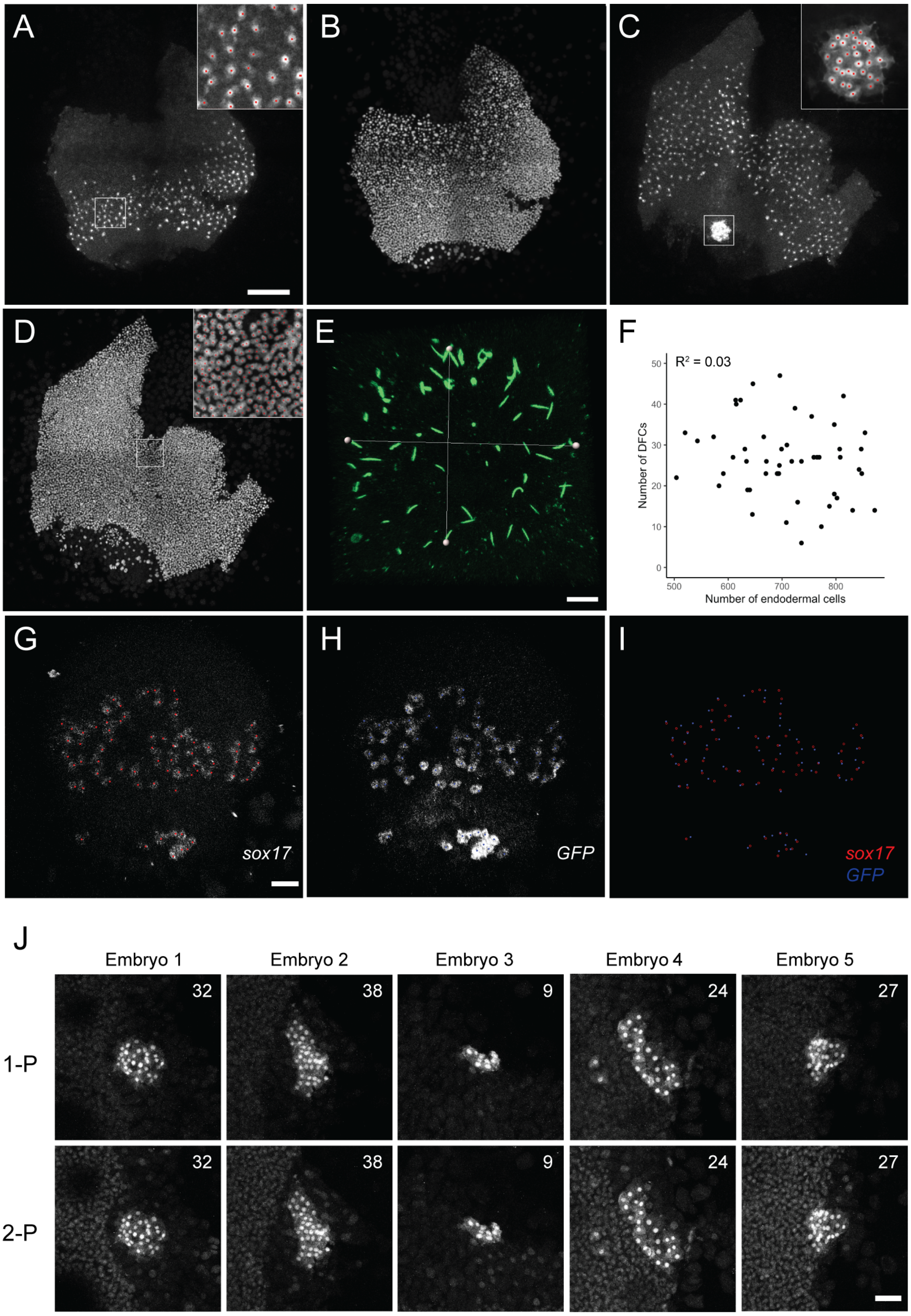


**Supplemental Figure 1, Related to Figure 1. Quantification of endodermal cells, DFCs, total cell number and KV cilia number on flat mount preparations.** (A-D) Confocal z-projections of a Tg[*sox17*:GFP] flat-mounted embryo at 75% epiboly stage. (A, B) ventral side; (C, D) dorsal side; (A, C) GFP immunodetection signal; (B, D) nuclei staining with Hoechst.

(A, C, D) Magnified regions show examples of cell quantification (red dots) for DFCs (taken separately, see Materials and Methods), endodermal cells and total cell number, respectively. Scale bar = 200 µm. (E) Confocal z-projection of an 8-somite embryo's KV, acetylated Tubulin (ac-Tub) immunodetection signal for individual cilia. The white lines indicate the two diameters used to calculate the area of the KV lumen as an ellipse. Scale bar = 10 µm. (F) Number of DFCs and endodermal cells detected per embryo, together with the coefficient of determination ($R^2$). (G-H) *sox17* and *GFP* transcript detection on the dorsal side of a Tg[*sox17*:GFP] embryo at 75% epiboly. Red and blue dots denote the area considered to be an individual cell, respectively; (I) Overlap image. Scale bar = 50 µm. (J) Comparison of DFC detection by 1-Photon (1-P) and 2-Photon (2-P) microscopy on five individual Tg[*sox17*:GFP] embryos at 75% epiboly. The number of DFCs detected is indicated at the top right corner of each image. Scale bar = 50 µm.

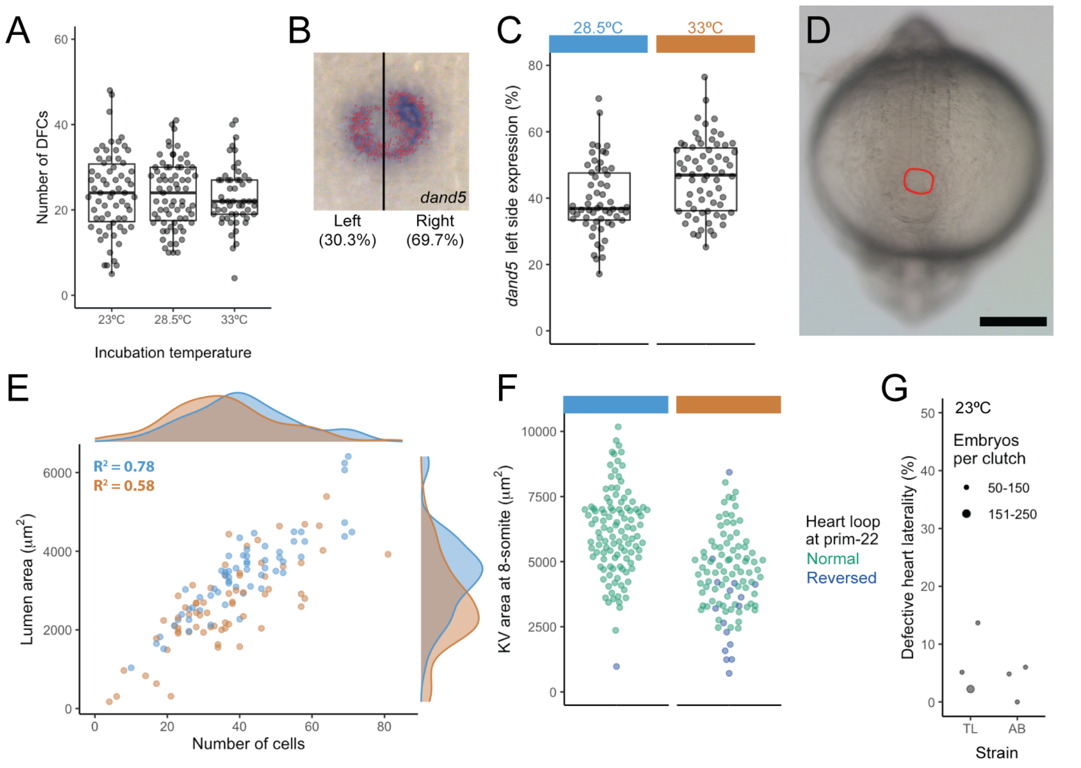


**Supplemental Figure 2, Related to Figure 2. Incubation temperature affects early LR signaling and KV size.** (A) DFC numbers at shield stage for different incubation temperatures (shield stage embryos, AB strain). (B) *dand5* expression in the KV of a 10-somite embryo detected by *in situ* hybridization. Relative levels of *dand5* expression on the left and right side as determined after color thresholding in Fiji. (C) Column scatter plot of the *dand5* left side expression in embryos incubated at 28.5ºC (light blue) and 33ºC (orange), p = 0.017. (D) Photograph of the posterior part of an 8-somite stage embryo, showing the KV (red circle) below the notochord. Scale bar = 200 µm. (E) KV lumen area and cell number in TL embryos incubated at 28.5ºC and 33ºC. The distributions for these two parameters are shown as marginal histograms at the top and right and the coefficient of correlation is shown for each condition. Lumen area p < 0.001, cell number p = 0.04. (F) Quantification of the KV area in live embryos at 8-somite stage together with the heart laterality phenotype observed on the same embryos at prim-22 , after incubation at 28.5ºC or 33ºC, p <0.001. (G) Heart laterality phenotypes observed at prim-22 stage in embryos incubated at 23ºC.

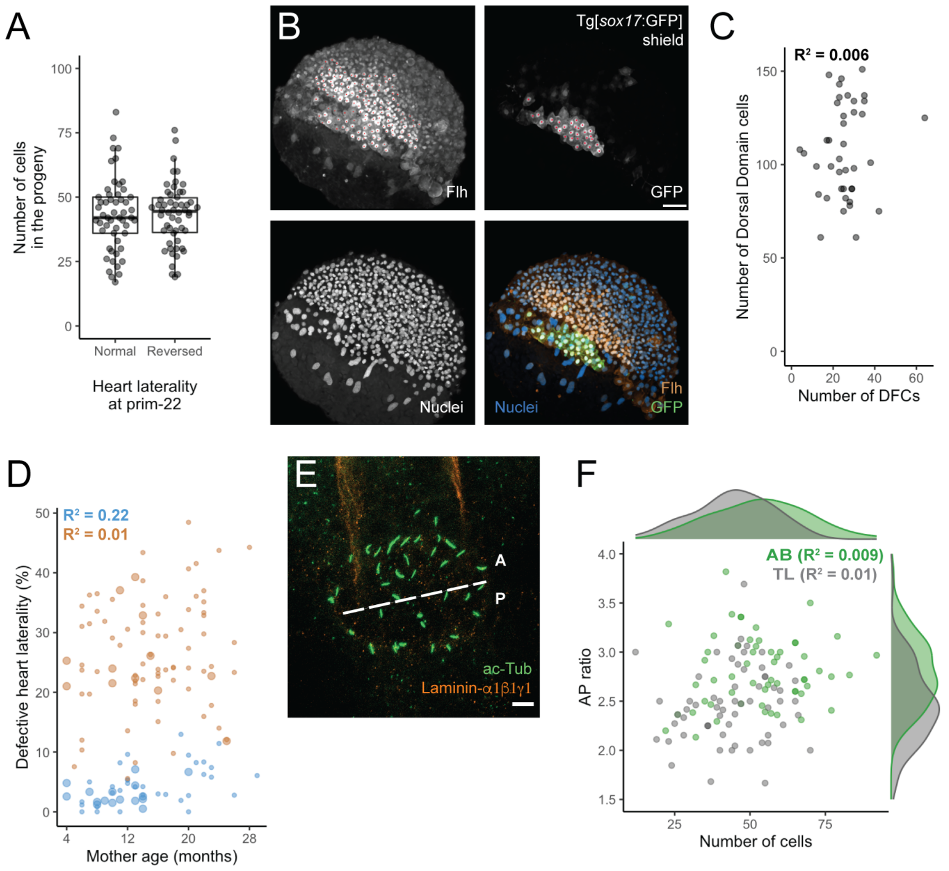


**Supplemental Figure 3, Related to Figure 3. Origin of DFC number fluctuations.** (A) Number of KV cells in the progeny of individuals that showed either normal or reversed heart laterality at prim-22 stage. (B) Confocal z-projections of the dorsal side of a Tg[*sox17*:GFP] shield stage embryo, positive cells are marked with a red dot for Flh and GFP immunodetection signal (upper panels). The lower panels display nuclei staining with Hoechst and a composite image of all three channels: nuclear staining: blue, Flh: orange, and GFP: green. Scale bar = 50µm (C) Correlation between the numbers of DFCs and dorsal domain cells detected per embryo. (D) Percentage of DHL versus the age of the mother used for each TL cross. (E) Confocal z-projection of a KV at 8-somite stage, ac-Tub (cilia) and Laminin-α1β1γ1 (notochord) immunodetection signals are shown in green and orange, respectively. The latter is used as reference to set the KV anterior and posterior regions and estimate the anteroposterior ratio. Scale bar = 10µm. (F) Antero-posterior ratio of cell numbers in the KV of AB and TL 8-somite embryos ($p < 0.001$) and the number of cells (same data as in Figure 3C for 8-somite stage). The distributions are shown as marginal histograms at the top and right, and the coefficient of correlation is shown for each strain.

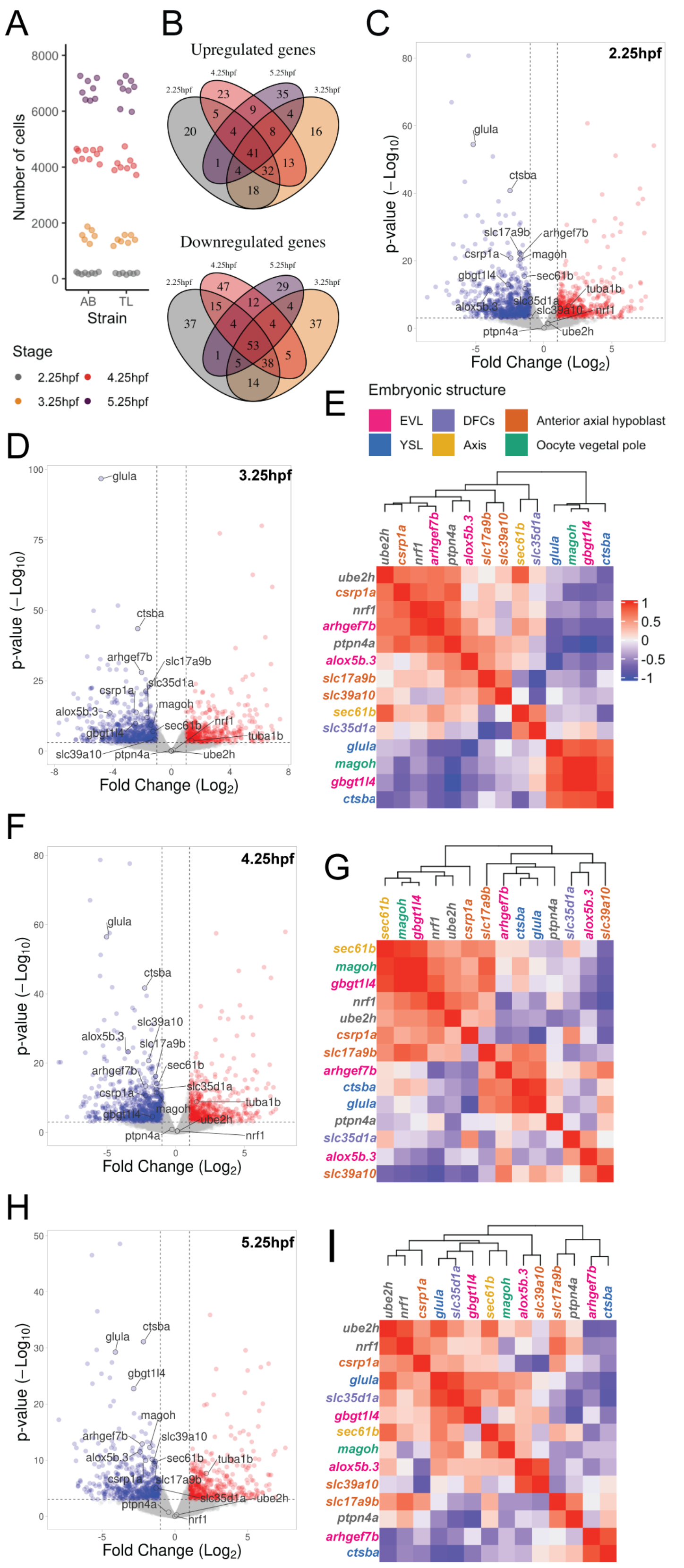


**Supplemental Figure 4, Related to Figure 3. Transcriptomic comparison of TL vs AB embryos during early embryogenesis.** (A) Total cell number estimation with 2-photon microscopy for embryos from the same batch used for transcriptomic analysis at each stage (color legend at the bottom). (B) Venn diagrams showing the intersection of the number of differentially expressed genes shared across different stages. (C, D, F, H) Volcano plots for TL/AB comparison at the timepoint indicated, with shared differentially expressed genes highlighted. (E, G, I) Heatmaps showing the pairwise Pearson correlation between the downregulated genes at 3.25, 4.25 and 5.25hpf, respectively, for TL embryos. (E) includes the color code for the genes by embryonic structure (outgroup genes in gray) and the scale for the correlations.
